# Adult porcine (*Sus scrofa*) derived inner ear cells possessing multipotent stem/progenitor cell characteristics in *in vitro* cultures

**DOI:** 10.1101/2021.01.22.427339

**Authors:** Printha Wijesinghe, Anand Sastry, Elizabeth Hui, Tristan A. Cogan, Boyuan Zheng, Germain Ho, Juzer Kakal, Desmond A. Nunez

## Abstract

The human inner ear compared with that of other mammalian species is very complex. Although the mouse’s cochlea is frequently studied the mouse’s inner ear continues to develop postnatally whilst the human inner ear is fully developed by the third month of gestation which leads one to question the applicability of findings based on research on mice to human regenerative therapies. Here, we report a novel *in vitro* culture of adult porcine (*Sus scrofa*) inner ear cells developed from post-mortem labyrinth specimens. Anatomical findings based on maximal transverse and vertical axial diameters and the length of the cochlear duct suggest that the pig’s cochlea is similar to the human cochlea. *In vitro* cultures of porcine cochlear and vestibular cells showed the persistence of both inner ear hair cell (HC), supporting cell (SC) and stem/progenitor cell characteristics across passages up to 6 based on scanning electron microscopy, fluorescence immunocytochemistry and quantitative reverse transcription polymerase chain reaction (RT-qPCR). Our findings showed that porcine cochlear and vestibular epithelia maintained multipotent stem/progenitor cell populations into adulthood although their regenerative capacities differed across the passages. The development of a viable and reproducible method to culture porcine inner ear cells provides an important investigative tool that can be utilized to study and evaluate the pathophysiological causes and cellular consequences of human inner ear disorders.

## Introduction

Human hearing and balance disorders are mostly attributed to damage to the mechanosensory hair cells (HCs) of the cochlear and vestibular sensory epithelia, respectively. These HCs are susceptible to a variety of insults including noise, ototoxic compounds, ageing and the interaction of adverse environmental and genetic factors [1]. Regardless of the cause, lost or damaged human cochlear HCs are never replaced, and the replacement of vestibular HCs occurs at levels too low to support significant functional recovery [2, 3]. In mammals, sensory HCs arise from embryonic progenitor cells during the embryonic period and in some species such as mice where the ear is not fully developed at birth, also during the early post-natal period [4]. There is as yet no evidence of newly generated *de-novo* mammalian auditory HCs in the mature cochlea [5].

Mammalian vestibular HC regeneration is believed to arise through nonmitotic trans-differentiation of supporting cells (SCs) [6, 7]. In contrast to mammals, non-mammalian vertebrates such as birds produce or regenerate auditory HCs after trauma, and thus can maintain optimum hearing function throughout their lives [8]. The observed patterns of cell division and differentiation in avian species suggest that their HCs regenerate from postembryonic progenitor cells [9]. The generation of new HCs from a renewable source of progenitor cells is a principal requirement for the development of an inner ear cell-based therapy [10].

The study of human HCs is severely limited because the cochlea is technically difficult to access, tissue harvest leads to severe and permanent hearing disability, and surgery involving inner ear tissue removal is comparatively rare. Hence other species are utilized in mammalian inner ear research. The mouse is the most commonly studied species due to its small size, high reproductive rate, large reported genetic database, variety of different strains, and the relatively low cost to procure and maintain study subjects. However, the mouse’s inner ear continues to develop postnatally whilst the human inner ear is fully developed by the third month of gestation [4] meaning that mice retain the capacity for auditory sensory HC regeneration after birth in contrast to humans which questions the applicability of findings based on research on mice to human regenerative therapies.

The pig has been considered as a superior model for the study of human diseases particularly for understanding complex conditions such as obesity, arthritis, cardiovascular, skin, and eye diseases [11]. Pigs have many similarities to human with respect to compatible organ size, immunology and physiological functions. Their high-quality annotated reference genome sequence and many known alleles presumed to cause diseases extend the potential of the pig as a biomedical study model [12, 13]. The morphology and the development of the inner ear of miniature pigs have been reported to be similar to that of humans [14].

Our objectives were to develop a viable method to isolate porcine inner ear cells from postmortem cochlear and vestibular epithelia, document the porcine inner ear anatomical structure, and report an *in vitro* mixed cell culture model that demonstrated the presence and persistence of inner ear HC, supporting cell (SC) and stem/progenitor cell characteristics.

## Materials and Methods

### Ethical Approval

This study was approved by the Biosafety Committee of the University of British Colombia (B14-0048, B18-0048), Vancouver, Canada and by the Animal Research Ethics Review, University of Bristol, Bristol, United Kingdom. All the experiments were performed in accordance with host institutional Policies and Procedures, Biosafety Practices and Public Health Agency of Canada guidelines as required.

### Tissue harvest, isolation and cell culture

Twenty-four temporal bone labyrinth capsules were harvested from 12 adult pigs (*Sus scrofa*) with ages ranging from 15-19 weeks. Six temporal bone labyrinths were selected at random for identification of the gross anatomy and histological study. The otic capsule of the temporal bones were isolated from euthanized adult pigs immediately post-mortem (within 2 hours) and placed in pre-chilled sterile Dulbecco’s phosphate-buffered saline (DPBS).

Four temporal bone labyrinths were used for *in vitro* cell cultures. The helicotrema of the cochlea was opened and the cochlear duct-extracted piecemeal. Vestibular tissue was harvested in part via the oval window, and by dissection of the bony semi-circular canals with special attention to harvesting darkly pigmented vestibular epithelium.

The harvested tissues were macerated under a dissection microscopic vision, keeping the cochlear and vestibular tissues separate. Each cochlear and vestibular tissues were collected in separate 15ml sterile tubes contaning 10ml DPBS and 2ml of 0.25% trypsin-EDTA and then incubated at 37^°^C for 20-25 minutes for trypsiniation. Tissues extracted from two temporal bone labyrinths were preserved separately in RNAlater, a protective reagent, and stored at -80^°^C for subsequent RNA extraction.

Following incubation, the tubes were centrifuged at 300 x g for 5minutes and the resulting cell pellets were re-suspended in 3ml of growth medium consisting of Dulbecco’s Modified Eagle Medium (DMEM) enriched with 10% Fetal Bovine Serum (FBS). About 500µl of the cellular suspension was then placed into each well of a 6-well cell culture plate pre-filled with 1.5ml of fresh growth medium.

The culture plate was placed into an incubator set at 37°C and 5%CO_2_. Half of the medium was replaced after a week taking care not to disturb the growing cells, and after a further 5-7 days all the medium was changed. When the cell growth reached >80% confluence (approximately 3 weeks), the cells were washed with DPBS buffer and trypsinized using 250µl of 0.25% trypsin-EDTA per well and incubated at 37^°^C for 5 minutes. Trypsinization was stopped by adding 9 ml of DMEM medium (without 10% FBS) and the pooled suspension was centrifuged at 300 x g for 5minutes. One-third portion of the cell pellet (passage 0 or P0) was re-suspended in fresh growth medium and placed in T_25_ vented culture flasks. The next generation of cells (passage 1 or P1 cells) was allowed to grow until >80% confluence was reached.

The cells were subsequently re-passaged again, up to P6, allowing for >80% confluence before each passage. The morphology of the cochlear and vestibular derived cells were recorded at each passage by a Phase-contrast Zeiss Axio Vert.A1 inverted microscope. Ultra-morphological features were obtained by a Scanning Electron Microscope (SEM) - Hitachi S4700 on passaged cells.

We have used House Ear Institute-Organ of Corti 1 (HEI-OC1) cells (gifted by Dr. F. Kalinec), a conditionally immortalized mouse auditory cell line as a positive control for cellular characterization studies [15]. HEI-OC1 cells were cultured under non-permissive conditions by incubating at 37^°^C and 5% CO_2_ in T_25_ vented culture flasks containing DMEM medium and 10% FBS without supplements.

#### Scanning electron microscopy on passaged inner ear cell cultures

Ultra-morphological features of P4 passaged porcine inner ear cells and HEI-OC1 cells were obtained by scanning electron microscopy. The porcine inner ear cells and HEI-OC1 cells were grown on poly-L-lysine coated cover slips under the above described culture conditions. At 100% confluence, cells were prepared for SEM [16] as follows: cells were washed using straight phosphate buffer (pH 7.4) and fixed for 30 minutes with 2.5% glutaraldehyde in 0.1 M sodium cacodylate buffer (pH 7.4), containing 2mM CaCl_2_. After fixation, cells were washed 3 times with 0.1M sodium cacodylate buffer (pH 7.4), each for 1 minute, post-fixed for 10-15 minutes with 1% osmium tetroxide (OsO4) in the same sodium cacodylate buffer and then washed. The cells were progressively dehydrated in a graded ethanol series, critical point-dried using CO_2_ and sputter-coated with gold/palladium (Au/Pd). The cells were then examined using a Hitachi S4700 SEM.

### Temporal bone sectioning, histology and inner ear anatomy

The bone labyrinths of six temporal bones were obtained as described above and fixed in 37% formaldehyde. The diameters of the six labyrinthine bones were measured with precision digital calipers. The bones were then decalcified by first immersing in 0.5M EDTA (pH 8.0) solution for 7 days. This was re-freshed with 0.5M EDTA (pH 8.0) solution for a minimum of four times over the 7 days. The bones were then immersed in 10% sucrose for 2 hours, followed by 20% sucrose for 24 hours and 30% sucrose for 24 hours. Four of the bones were then placed in an optimum cutting temperature (OCT) medium at room temperature.

A round window cochleostomy was fashioned under microscopic control in two of the cochleas. The length of the cochlea was determined by observing how far the active electrode array of a standard Nucleus 22 cochlear implant could be inserted via the round window.

All of the decalcified specimens embedded in OCT were frozen in isopentane held in the vapour phase of liquid nitrogen. Serial 5μm sections of the bones were cut with a microtome and placed onto Superfrost Gold slides. Sections were air dried for 48 hours and fixed in acetone prior to staining with haematoxylin and eosin or haematoxylin/van Gieson stains.

### Cellular characterizations

#### Fluorescence immunocytochemistry (ICC)

At passages 0, 1 and 4, the cochlear and vestibular cell cultures were examined for the expression of the inner ear HC markers (myosin VIIa and prestin), SC markers (cytokeratin 18 and vimentin) and multipotent stem/progenitor cell markers (nestin and Sox2). Cells grown on 8-well chamber slides (Thermo Scientific Nunc; Lab Tek) were immunostained at confluence ≥80%. Initially, culture medium was removed and the cells were washed 3 times in DPBS, each wash was of 1 minute duration. The cells were then fixed by incubation in 4% paraformaldehyde for 15 minutes, followed by permeabilization in 0.1% Triton-X 100 for 15 minutes. Thereafter, the cells were blocked using 3% Bovine Serum Albumin (BSA) at room temperature for 30 minutes prior to incubation at 4°C overnight with primary antibodies [myosin VIIa (inner HC marker) 1:100 dilution (rabbit polyclonal-ab3481, ABCAM); prestin (outer HC marker) 1:100 dilution (goat polyclonal-SC22692, Santa Cruz Biotechnology); nestin (stem/progenitor cell marker) 1:100 dilution (rabbit polyclonal-ab92391, ABCAM); Sox2 (stem/progenitor cell marker) 1:100 dilution (rabbit polyclonal-ab97959 ABCAM); cytokeratin-18 (an epithelial cell marker) 1:50 dilution (mouse monoclonal - ab668, ABCAM) and vimentin (mesenchymal cell marker) 1:200 dilution (mouse monoclonal-ab20346, ABCAM and rabbit polyclonal-PA5-27231, Invitrogen)] dissolved in 3% BSA.

The following day, primary antibodies were drained and the chamber slides washed 3 times, each for 1 minute, in DPBS. Then, the cells were incubated at room temperature with secondary antibodies in the dark [goat anti-rabbit alexa fluor®488 1:500 dilution (ab150113, ABCAM); goat anti-mouse alexa fluor®488 1:500 dilution (ab150077, ABCAM); donkey anti-rabbit alexa fluor®488 1:500 dilution (A21206, Invitrogen); donkey anti-mouse alexa fluor®568 1:500 dilution (A10037, Invitrogen); and donkey anti-goat alexa fluor®488 1:500 dilution (A11055, Invitrogen)], respectively to the primary antibodies and shaken gently for 1 hour. The cells were then mounted with ProLong™ Gold Antifade Mountant with DAPI (P36931, Invitrogen). Images were captured using a Zeiss Axio Vert.A1 Inverted Microscope. HEI-OC1 cells served as the positive controls, and cells treated without primary antibody served as the negative controls.

### Quantitative reverse transcription polymerase chain reaction (RT-qPCR) on adult porcine derived cochlear and vestibular cells

RNA was extracted from tissues and primary cell cultures of adult porcine derived cochlear and vestibular cells that were grown on T_25_ culture flasks. The level of mRNA expression of target genes *myosin VIIa, prestin, nestin, Sox2, cytokeratin 18* and *vimentin* was determined at the tissue level and at passages 0, 2, 4 and 6 using the comparative Cycle threshold (ΔΔCt method). A housekeeping gene for these experiments was selected by testing three candidate genes *glyceraldehyde-3-phosphate dehydrogenase* (*Gapdh*), *beta-actin* (*b-Act*) and *hypoxanthine phosphoribosyltransferase 1* (*Hprt1*) simultaneously. RefFinder, a web-based tool was used to select the most stable of the candidate housekeeping genes tested by computing the weighted geometric means of their individual rankings derived by four widely used methods [17]. Target genes were also determined in positive control HEI-OC1 cells.

Primer3Plus software [18] was used to design the forward and reverse primers for both porcine inner ear samples and HEI-OC1 cells (S1 Table and S2 Table, respectively). Primers were designed for the genes associated with porcine inner ear HCs, SCs and stem/progenitor cell characteristics using the GenBank (NCBI) database.

RNAs (RNeasy® mini kit, QIAGEN) were extracted from porcine cochlear and vestibular tissues preserved in RNAlater held at -80°C and from porcine (at P0, P2, P4 and P6) and HEI-OC1 derived cell culture pellets which were dissolved in 350µl RLT buffer containing 0.01% 14.3M β-mercaptoethanol according to the manufacturer’s instruction. The quantity and quality of extracted RNAs were determined prior to cDNA preparation. We performed cDNA synthesis with SuperScript™ VILO™ cDNA Synthesis Kit (Invitrogen) under the following reaction conditions: 25^°^C for 10 minutes, 42^°^C for 60 minutes and 85^°^C for 5 minutes (in a BioRadT100™ Thermal Cycler).

Synthesized cDNAs were then diluted to a concentration of 5ng/µl for RT-qPCR in a StepOnePlus™ Instrument (Applied Biosystems) using 96-well plates and a SYBR Select Master Mix reagent. In brief, the RT-qPCR reaction mix per well consisted of 1µl of HyPure™ Molecular Biology Grade Water, 5µl SYBR Select Master Mix at the manufacturer’s supplied concentration, 1µl of each forward and reverse primer (10µM) and 2µl of diluted cDNA (5ng/µl). After the reaction mix was added to the wells, the plate was centrifuged for few a seconds in a Mini PCR Plate Spinner. A RT-PCR consisted of an initial denaturing step of 95^°^C for 10 minutes and followed by 40 amplification cycles of 15 seconds at 95^°^C and 1minute at 60^°^C for each cycle. Three replicates were taken from each sample.

Relative mRNA levels were determined using the comparative cycle threshold method at a cut off of Ct <35. The relative mRNA levels were expressed as the mRNA copies of the genes of interest per 1000 copies of *Hprt1* mRNA [2^-ΔCt^/1000 = 1000/2^ΔCt^= 1000/2^^(avg.target gene Ct − avg. housekeeping gene Ct)^] [19, 20].

### Statistical Analysis

The differences in the normalized mean Ct values of target genes between porcine cochlear and vestibular cells, and between porcine cochlear and HEI-OC1 cells were analyzed statistically using Student’s *t*-test at a Benjamini-Hochberg corrected significance level of p ≤0.05. The strength of correlation among different target gene expression levels within porcine cochlear and vestibular cell cultures was analyzed using a non-parametric Spearman’s rank correlation coefficient test. Analyses were conducted using SPSS version 25.0 (IBM Corp., Armonk, New York). The level of statistical significance was set at p <0.01.

## Results

### Adult porcine inner ear anatomy

Based on six temporal bone labyrinths selected at random from 12 adult pigs, the gross anatomy of the porcine labyrinth showed some similarity to human’s. They both have a recognizable cochlear spiral though the pig’s consisted of 3.5 turns compared to the human’s 2.5 turns. There was a separate vestibular compartment arranged in raised bone canals. The six cochleas were found to have a mean maximal diameter and height of 7.99 and 3.77mm, respectively (Table 1). Average values ± standard error of mean are given. “A” denotes maximum axial diameter measured from the round window niche to the most lateral part of the basal turn of the cochlea. “B” denotes vertical axial diameter measured from helicotrema to the basal turn of the cochlea. “C” denotes short axis diameter measured between two opposite sides of the basal turn of the cochlea at right angles to measurement A. Measurements were taken using precision digital calipers to the nearest millimeter. Fig 1A illustrates how these measurements were made on a demineralised labyrinth.

**Table 1:** Summary of gross measurements of six porcine inner ear specimens.

| Cochlea | A | B | C |
| --- | --- | --- | --- |
| 1 | 7.97 | 3.72 | 7.38 |
| 2 | 8.01 | 3.66 | 7.31 |
| 3 | 8.03 | 3.70 | 7.32 |
| 4 | 7.97 | 3.95 | 7.40 |
| 5 | 7.96 | 3.76 | 7.36 |
| 6 | 7.98 | 3.85 | 7.37 |
| Average ±<br>standard error of<br>mean | 7.99 ± 0.01 mm | 3.77 ± 0.04 mm | 7.36 ± 0.01 mm |

**Fig 1.**
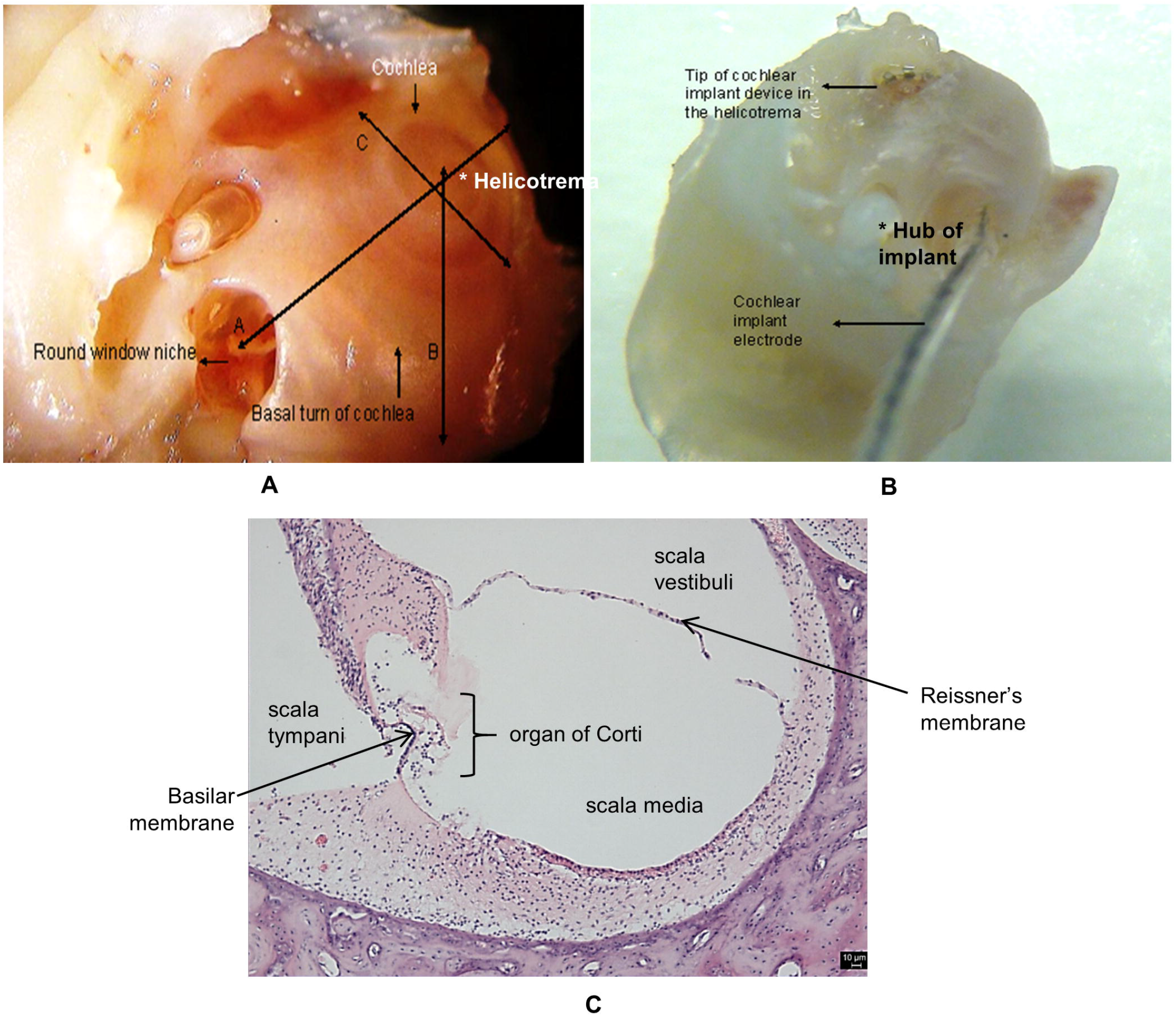
Adult porcine inner ear anatomy. **A)** A low power (10X) digital photomicrograph illustrates areas measured on a demineralized porcine inner ear that revealed a cochlea with three and half turns. Measurements were made as follows (in mm): A-Maximum axial diameter-measured from round window niche to the most lateral part of basal turn of the cochlea; B-Vertical axial diameter-measured from helicotrema to basal turn of cochlea; C-Short axis diameter-measured between two opposite sides of the cochlea basal turn at right angles to A. **B)** A low power (10X) digital photomicrograph of a demineralized porcine inner ear showing the cochlea and the position of the cochlear implant electrode array. A cochlear implant electrode array has been inserted through a round window cochleaostomy to determine the cochlea length. The tip of the implant array can be seen at the helicotrema and the hub at the level of the round window. **C)** Photomicrograph of a haematoxylin and eosin stained mid-modiolar section through the *Sus scrofa* cochlea. Cross sectioning of the *S. scrofa* cochlea demonstrates compartments consistent with the scala tympani, media and vestibuli partitioned by the basilar and Reissner’s membranes (indicated by arrows) identical to the arrangements found in the human cochlea.

Full insertion of a Nucleus C22 cochlear implant active electrode assembly through a round window cochleostomy was performed in two specimens such that the marker hub was level with the bony margin of the round window (Fig 1B). The tip of the electrode could be seen at the apex of the decalcified labyrinth in each case, leading to the conclusion that the pig’s cochlear duct length is within 25.5-35.1 mm range. A Haematoxylin and Eosin stained cross-section shows the pig’s bony cochlea to be partitioned into three compartments consistent with the scala tympani, media and vestibuli separated by the basilar and Reissner’s membranes identical to the arrangements in human inner ear (Fig 1C). The cochlear duct contained abundant tissue for cell culture.

### Cellular characterizations

#### Morphological characteristics of inner ear cells in in vitro culture

Porcine inner ear cells started to grow within 7-10 days of plating under *in vitro* conditions. Phase contrast microscopy illustrated that poricne inner ear derived cells at P0 contained a mixture of spindle shaped and flattened polyhedral shaped cells but by P4 there was a predominance of spindle shaped cells (Fig 2A to 2D). Proliferating cochlear tissue at P0 and P4 consisted of cell islands as shown in Fig 2A and 2B, respectively. Cells of similar appearance were seen in proliferating vestibular tissue at P0 and P4 (Fig 2C and 2D, respectively). Primary cell cultures consisted of heterogenous cells and formed monolayers which showed a high level of adhesion onto plastic surfaces. HEI-OC1 cells predominantly consisted of spindle shaped cells (Fig 2E) similar to P4 and later porcine derived inner ear cultured cells.

**Fig 2.**
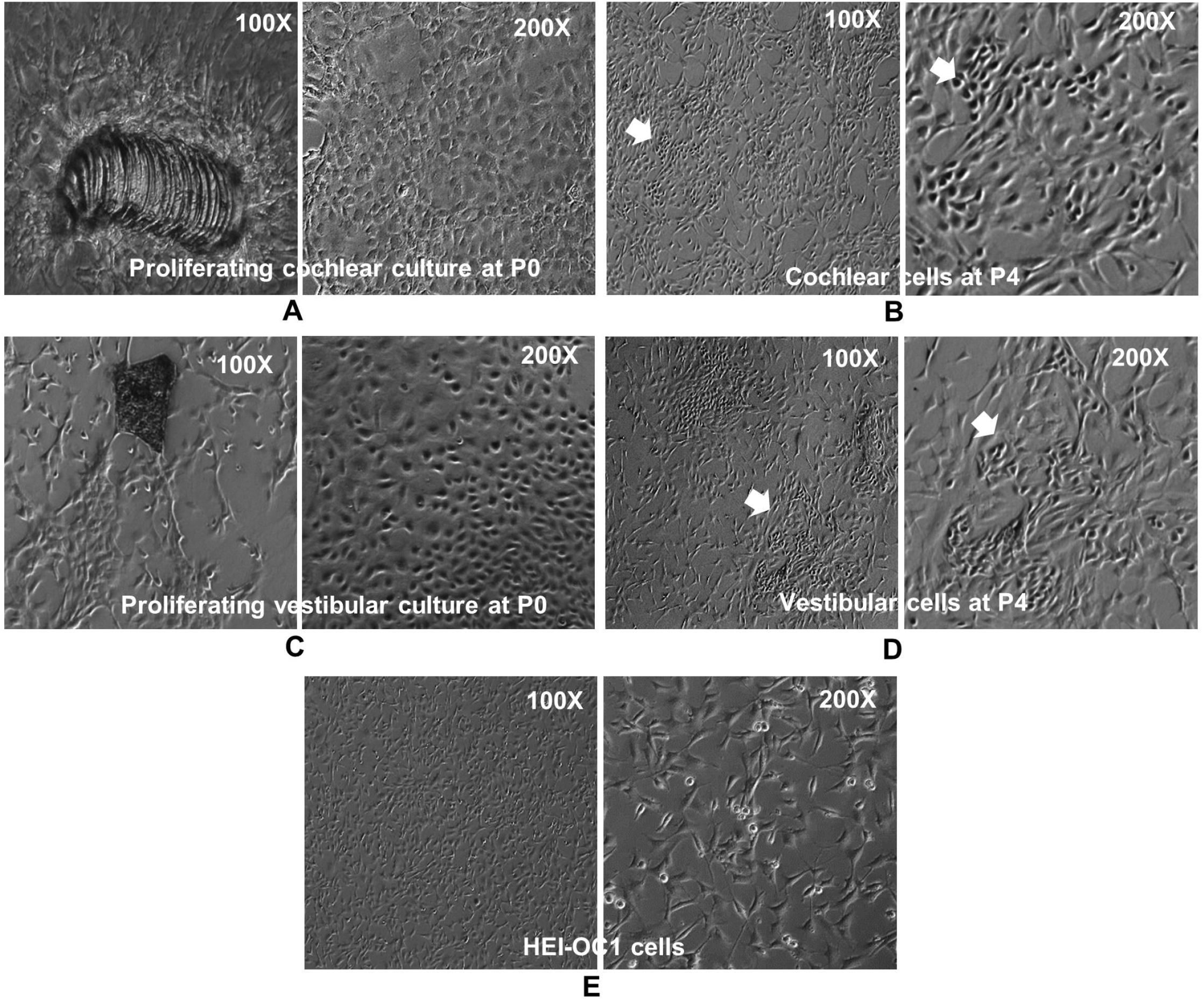
Morphology of inner ear *in vitro* cell cultures demonstrated by phase contrast microscopy. **A)** and **C)** Proliferating cochlear and vestibular cultures at passage 0; **B)** and **D)** Cochlear and vestibular cell cultures consisted of cell islands at passage 4 (indicated by white arrows); **E)** Positive control HEI-OC1 cells. The magnifications are given in each panel.

Porcine inner ear cells and HEI-OC1 cells grown on coated 8-well chamber slides showed similar sphere-forming charactericitics *in vitro* using phase-contrast microscopy (Fig 3A). SEM revealed that the sphere-forming HEI-OC1 (Fig 3B) porcine cochlear (Fig 3C and porcine vestibular cells (Fig 3D) were oblong in shape with large and small cytoplasmic protrusions on their surfaces.

**Fig 3.**
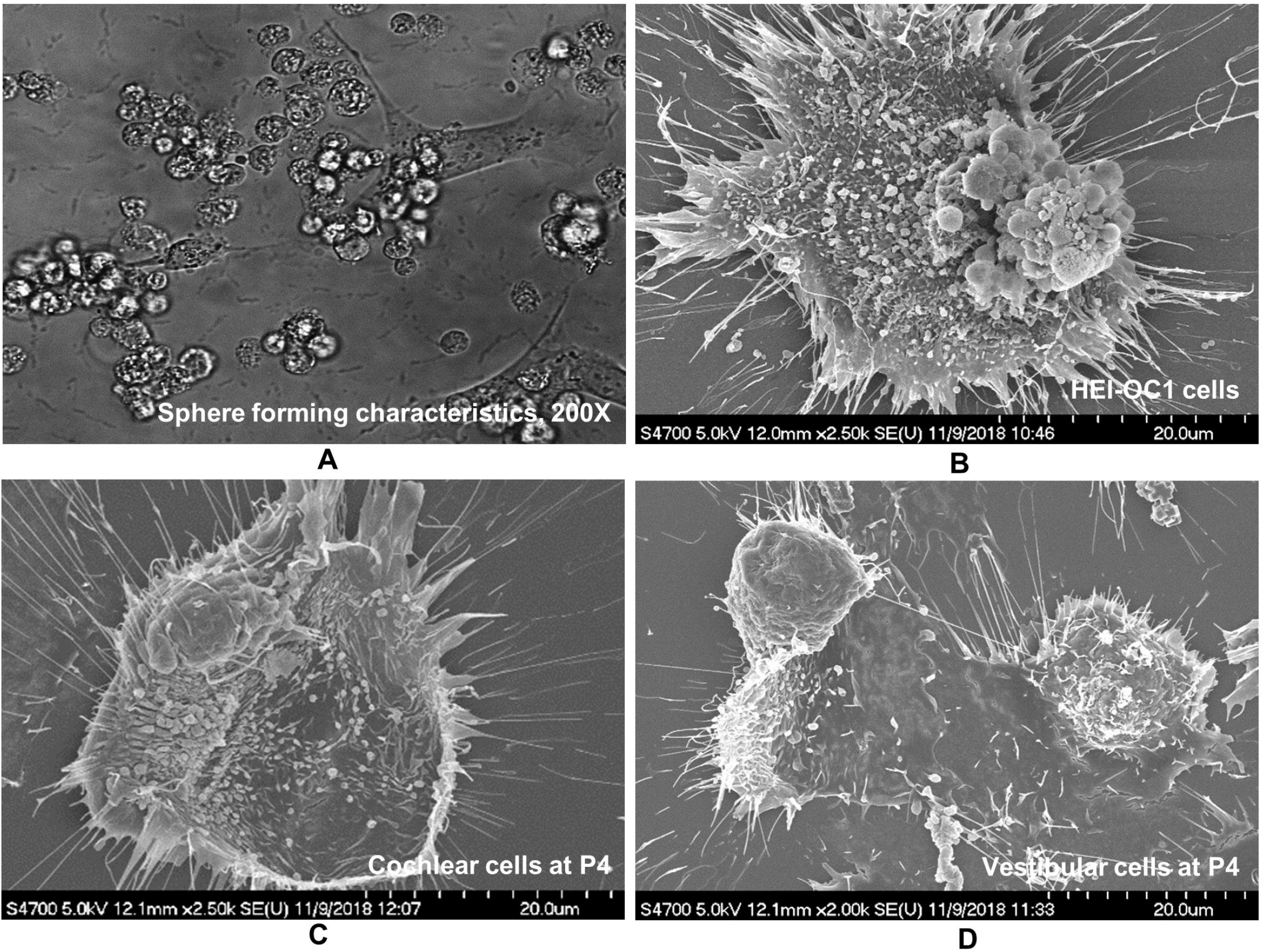
Sphere-forming characteristics of inner ear cells in *in vitro* cultures. **A)** Phase contrast microscopy demonstrated sphere-forming characteristics of porcine inner ear cells and HEI-OC1 cells grown on coated 8-well chamber slides. **B)** SEM revealed sphere-forming cells within the HEI-OC1 cultures that demonstrated prominent surface protrutions and cytoplasmic projections. **C)** Sphere-forming cells in porcine cochlear cultures similarly bore considerable protrusions on their surfaces and cytoplasmic projections. **D)** Sphere-forming cells in porcine vestibular cultures were similar though with less protrusions on their surfaces. Passaged 4 porcine inner ear *in vitro* cultures were used for SEM study.

Higher resolution SEM images of porcine derived inner ear cells revealed sphereical structures arising from cells with ciliary-type surface projections (Fig 4) *in vitro*. Cells with ciliary-type projections in HEI-OC1 culture (Fig 4A) and in porcine cochlear culture at P4 (Fig 4B) showed disorganized stereocilia-like structures with appendages suggestive of broken inter-stereociliary links (indicated by black arrows). Sphere-forming cells in the porcine vestibular cultures at P4 bore microvillar-like projections that varied in length some long and others short (Fig 4C).

**Fig 4.**
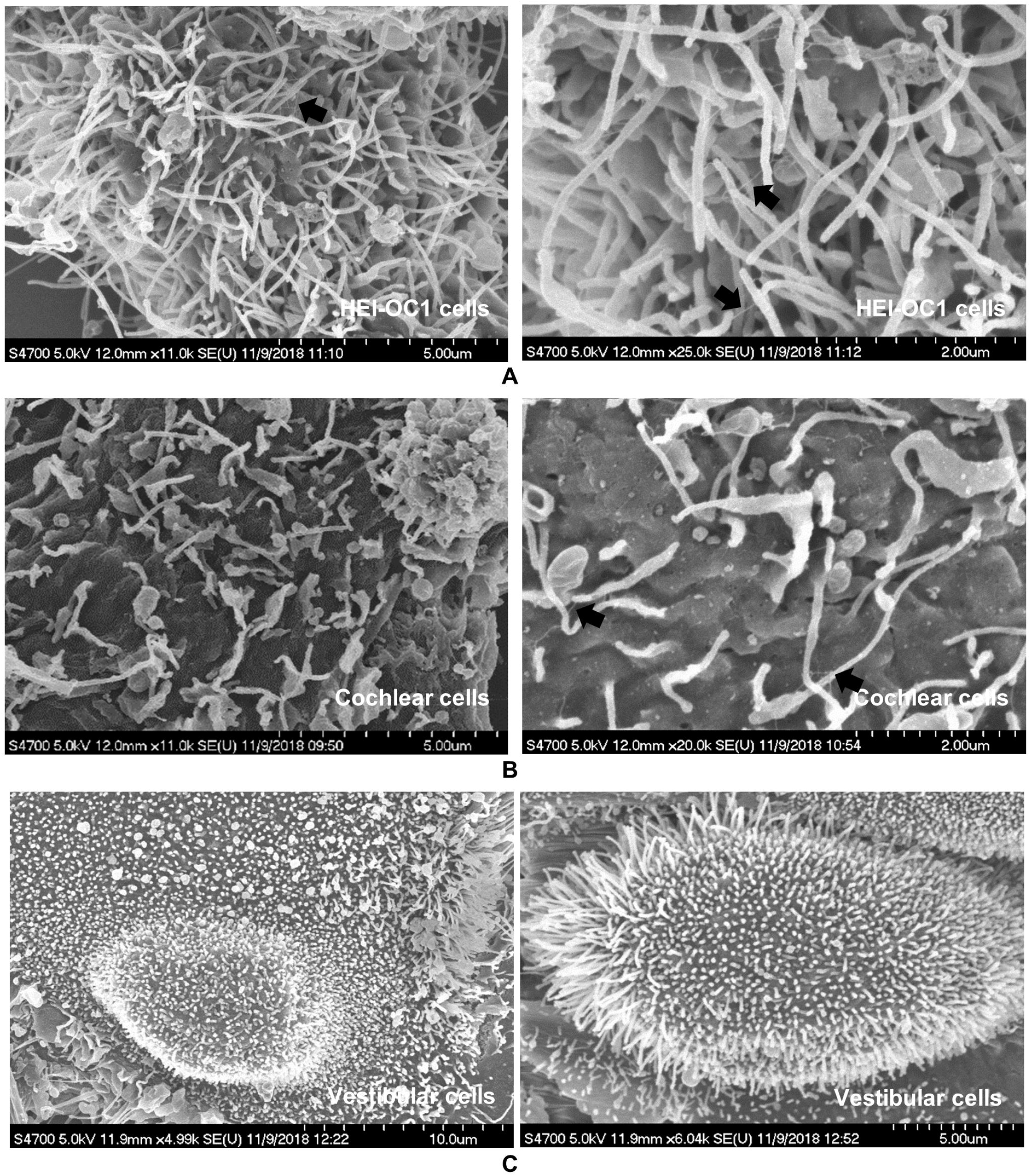
SEM characteristics of cells bearing ciliary-type projections on their surfaces. **A)** and **B)** Higher resolution SEM of sphere-forming cells with disorganized stereocilia-like structures with broken inter-stereociliary links (indicated by black arrows) in HEI-OC1 and porcine cochlear cells, respectively. **C)** Porcine vestibular cells are similarly shown to possess microvillar-like projections on their surfaces that vary in length though lacking any directional arrangement into rows of increasing length. Passage 4 porcine inner ear *in vitro* cultures were used for SEM study. Image resolution is indicated in each panel in micrometers.

#### Fluorescence immunocytochemistry

Stem/progenitor cell markers nestin and Sox2 were identified in adult derived porcine inner ear *in vitro* cultures (Fig 5). Nestin positive cells were plentiful in both cochlear (Fig 5A) and vestibular (Fig 5G) cell cultures at P1 and P4, respectively. Sox2 was localized in the nuclei of few porcine cochlear (Fig 5C) and vestibular (Fig 5H) cells at P1 and P4, respectively. More cells in both cochlear and vestibular cultures were positive for nestin rather than for Sox2 across the passages. Cell islands found in porcine vestibular cultures elicited strong signals to both nestin and Sox2 markers (indicated by white arrows). HEI-OC1 cells demonstrated a similar nestin (Fig 5B) and Sox2 (Fig 5D) expression pattern to porcine cochlear cells. Additionally, some of the globular or spherical shaped cells in porcine inner ear and HEI-OC1 cell cultures expressed both nestin and Sox2 (indicated by yellow arrows) markers. SEM demonstrated possible corresponding globular or spherical shaped cells with protrusions and long villi on their surface in porcine cochlear and HEI-OC1 cell cultures as shown in Fig 5E and 5F, respectively. These types of cells were observed consistently across all passages.

**Fig 5.**
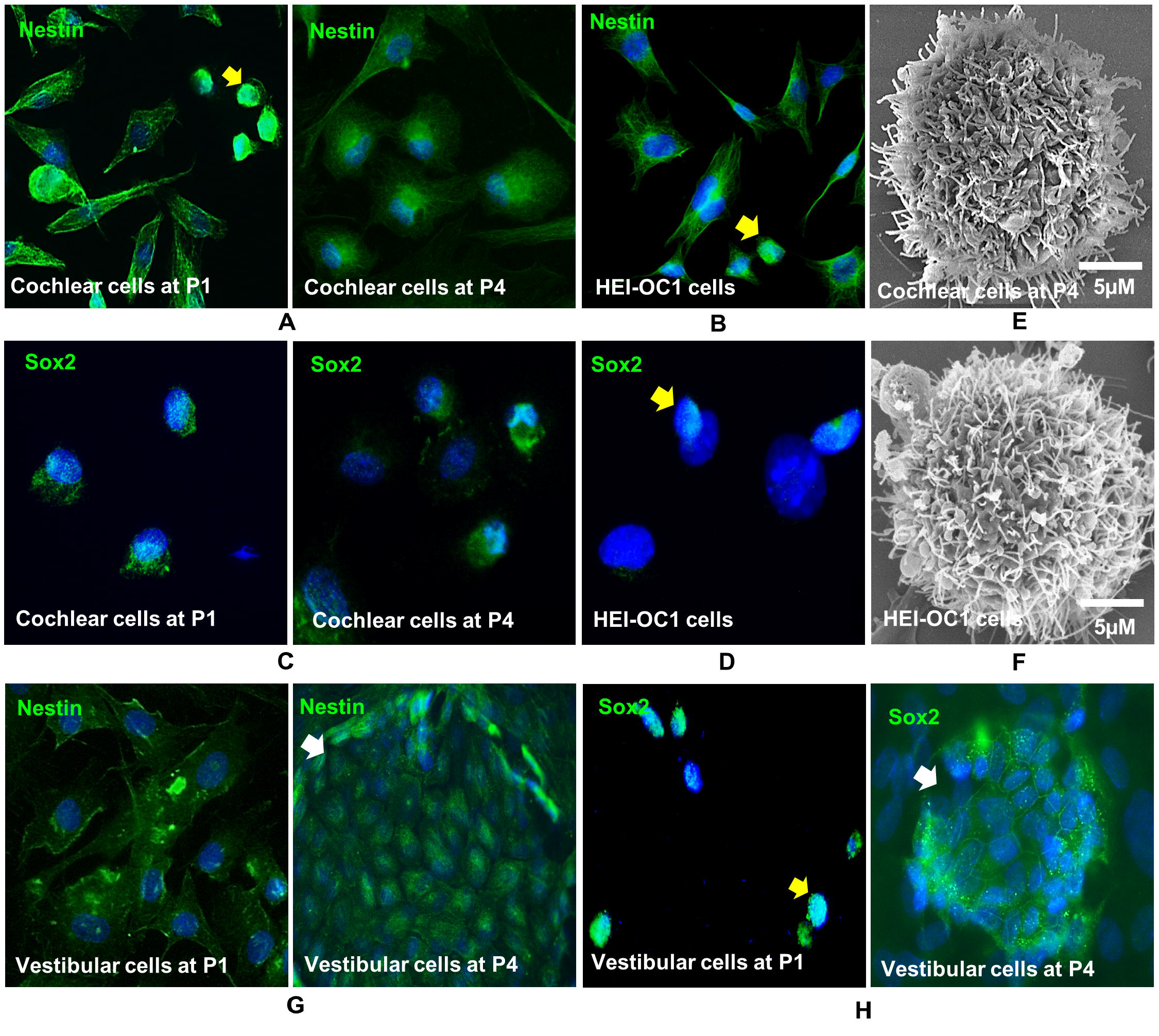
Presence of multipotent stem/progenitor cells in *in vitro* cultures. **A)** and **G)** Nestin positive cells were identified in the cochlear and vestibular cells at passages 1 and 4, respectively. **C)** and **H)** Sox2 positive nuclei were detected in the cochlear and vestibular cells at passages 1 and 4, respectively. **B)** Nestin positive cells in HEI-OC1 cells; **D)** Sox2 positive nuclei within the HEI-OC1 cells; **E)** and **F)** SEM images of a globular or spherical shaped cell with protrusions and long villi on its surface identified within the porcine cochlear and HEI-OC1 cell cutures, respectively. White arrows indicate the cell islands positive to both nestin and Sox2 markers within vestibular cultures at passage 4. DAPI was used to stain the nuclei (blue). Phase contrast microscopic images are given at magnification 400X. SEM image resolution is indicated in each panel in micrometers.

HC markers myosin VIIa and prestin positive cells were identified in porcine derived inner ear cell cultures based on immunofluorescence staining, although the intensity and dispersion varied across the cell passages (Fig 6).

**Fig 6.**
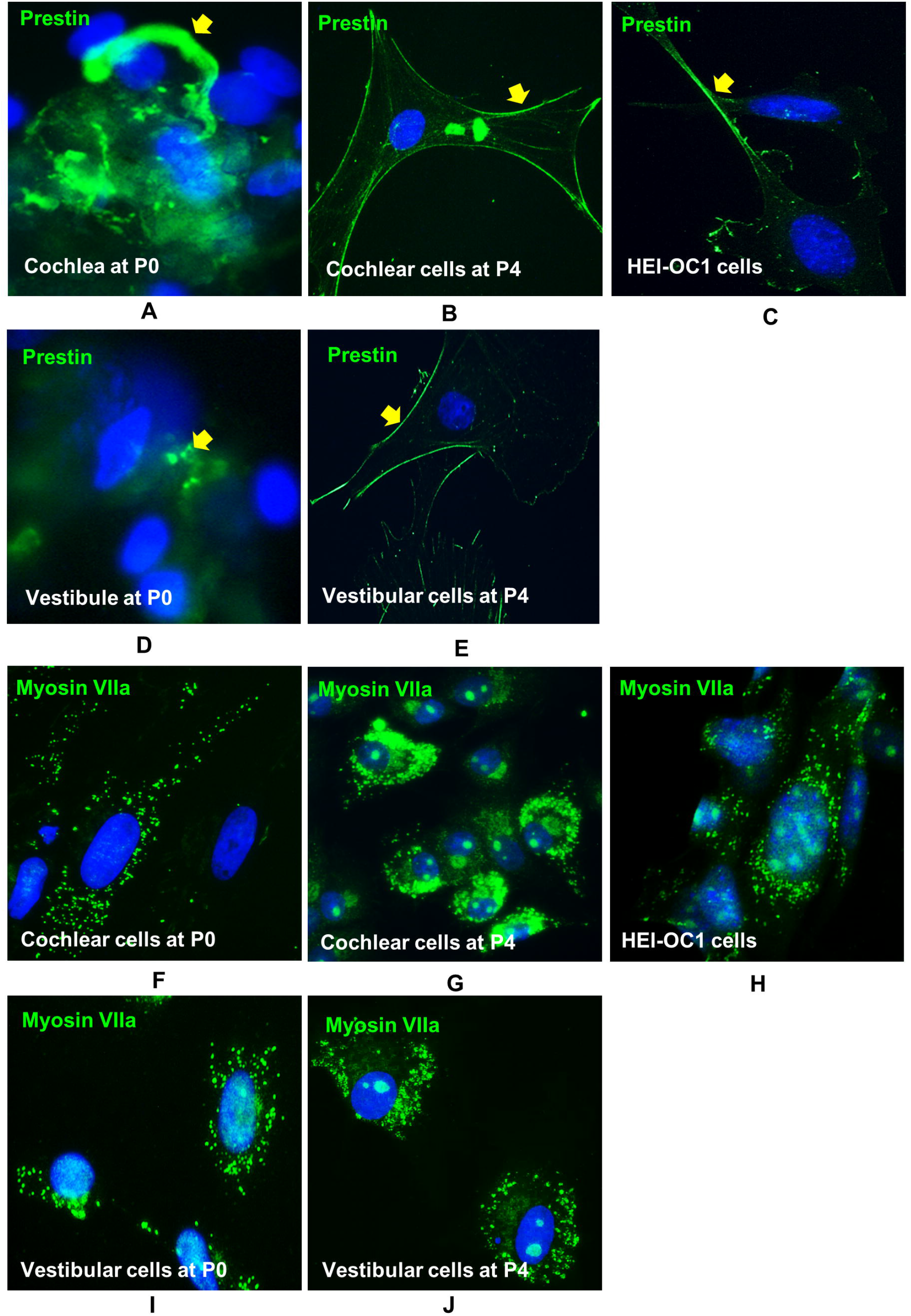
Localization of hair cell markers prestin and myosin VIIa. **A)** and **D)** Prestin localization identified in proliferating cochlear and vestibular tissues, respectively at passage 0 (indicated by yellow arrows). **B)** and **E)** Prestin plasma membrane localization strongly detected in cochlear and vestibular cells at passage 4 (indicated by yellow arrows). **C)** Prestin expression identified in HEI-OC1 cells. **F)** and **G)** Myosin VIIa expressions at passages 0 and 4 identified in cochlear cell cultures. **I)** and **J)** Myosin VIIa expressions at passages 0 and 4 identified in vestibular cell cultures. **G)** Myosin VIIa expression identified in HEI-OC1 cells. DAPI was used to stain the nuclei (blue). Phase contrast microscopic images are given at magnification 400X.

Prestin was localized in proliferating tissues of both the cochlea (Fig 6A) and vestibule (Fig 6D) at P0 and the expression was visibly stronger in cochlear than in vestibular *in vitro* cultures (indicated by yellow arrows). At P4, prestin expression was localized to the plasma membrane of cochlear (Fig 6B) and vestibular cells (Fig 6E) as indicated by yellow arrows. Prestin expression in HEI-OC1 cells was similarly localized as shown in Fig 6C. Prestin expression levels in both adult porcine derived cochlear and vestibular cells were visibly stronger than that of the positive control HEI-OC1 cells.

Myosin VIIa expression was strong and localized within the cytoplasm in adult porcine derived cochlear (Fig 6F and 6G) and vestibular (Fig 6I and 6J) cultured cells and in HEI-OC1 cells (Fig 6H). In contrast to the prestin marker, at P0, myosin VIIa expression was widely distributed throughout the porcine cochlear and vestibular derived *in vitro* cultured cells in keeping with diffuse cytoplasmic apical projections (Fig 6F and 6I, respectively). At P4, the myosin VIIa expression levels in porcine cochlear and vestibular cells (Fig 6G and 6J, respectively) were visibly stronger than in HEI-OC1 cells (Fig 6H) or P0 porcine derived cells. At P4, myosin VIIa positive signals were densely packed in both the porcine cochlear and vestibular cells compared to P0 cells.

At P0, SC protein markers, particularly cytokeratin 18 was vigorously expressed compared to vimentin in both cochlear (Fig 7A and 7B) and vestibular (Fig 7B and 7D) cultures, respectively. Cytokerain 18 expressing cells were compartively larger in size and polyhedral in shape. Vimentin expressing cells were comparatively smaller in size and mostly spindle shaped. At P4, vimentin was comparatively strongly expressed in cochlear and vestibular cells (Fig 7E and 7F, respectively), in contrast, cytokeratin 18 was weakly expressed in both cochlear and vestibular cells (Fig 7G and 7H, respectively). Vimentin and cytokeratin 18 protein experssions were comparable to each other in HEI-OC1 cells (Fig 7I and 7J, respectively) and they were smaller in size than porcine cells and mostly spindle shaped.

**Fig 7.**
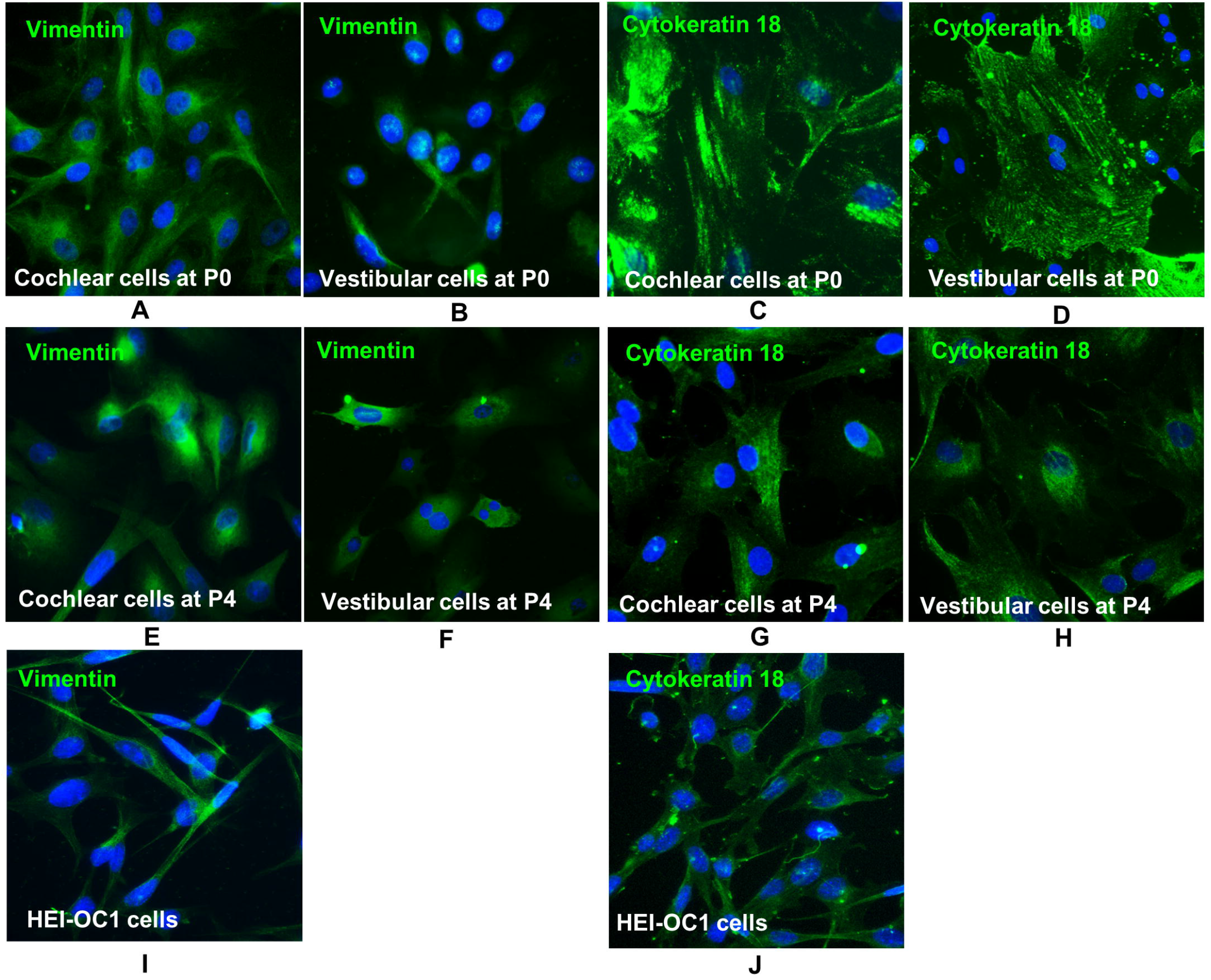
Immunolocalization of supporting cell markers vimentin and cytokeratin 18. Expression for: **A)** and **B)** vimentin at passage 0 in cochlear and vestibular cultures, respectively; **C)** and **D)** cytokeratin 18 at passage 0 in cochlear and vestibular cultures, respectively; **E)** and **F)** vimentin at passage 4 in cochlear and vestibular cultures, respectively; **G)** and **H)** cytokeratin 18 at passage 4 in cochlear and vestibular cultures, respectively; **I)** and **J)** vimentin and cytokeratin 18 positive HEI-OC1 cells, respectively. DAPI was used to stain the nuclei (blue). Phase contrast microscopic images are given at magnification 400X.

Double antibody labelling for myosin VIIa and cytokeratin 18 elicited strong signals within the porcine inner ear and HEI-OC1 cell cultures (Fig 8). Porcine vestibular cultures contained double-labelled epithelial cell islands (Fig 8A) at P4. Similarly, porcine cochlear culture showed double-labelled cells (Fig 8B) at P4, and the expression patterns were comparable to HEI-OC1 cells (Fig 8C).

**Fig 8:**
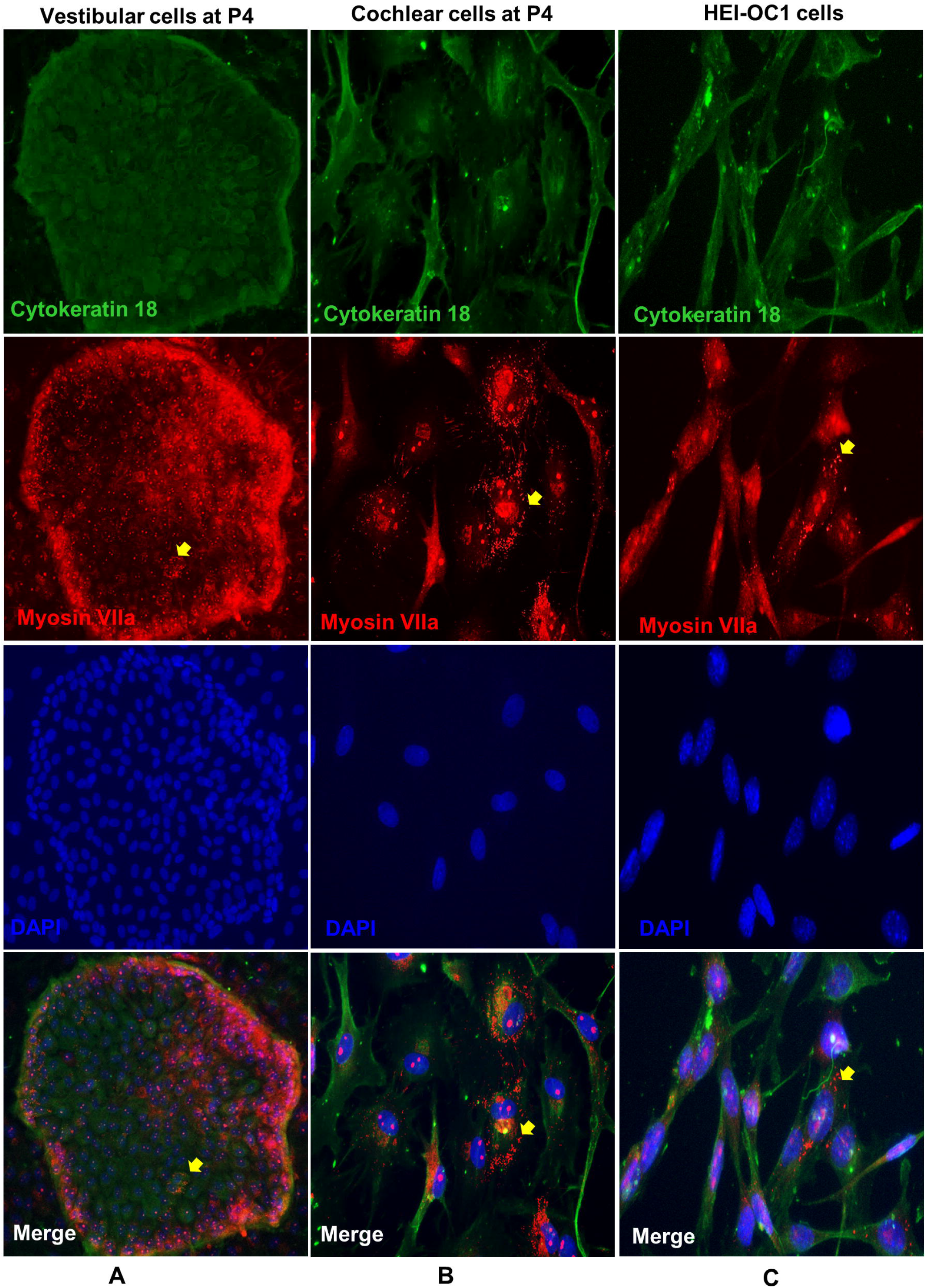
Double-labelled immunofluorescence demonstrated the co-localization of inner ear HC and SC markers. **A)** and **B)** Porcine vestibular and cochlear cells demonstrated dual labelling for antibodies cytokeratin 18 (green) and myosin VIIa (red) at passage 4; **C)** Double-labelled cells for cytokeratin 18 and myosin VIIa markers in HEI-OC1 cells. Closely packed myosin VIIa positive apical projections around the nucleus are indicted by yellow arrows. DAPI was used to stain the nuclei (blue). Phase contrast microscopic images are given at magnification 400X.

#### Relative gene expressions

In this study, three housekeeping genes, *b-actin, Hprt1* and *Gapdh*, were tested to determine RNA expression stability for RT-qPCR data normalization in cochlear and vestibular cell cultures throughout the passages. In cochlear cell cultures, from harvested tissues to P6, mean Ct value ± standard deviation (S.D.) for *Hprt1* was 23.41 ± 1.01, *Gapdh* was 16.59 ± 1.63, and *b-Act* was 14.38 ± 2.18. In vestibular cell cultures, from harvested tissues to P6, mean Ct value ± S.D. for *Hprt1* was 22.4 ± 1.01, *Gapdh* was 15.71 ± 1.58, and *b-Act* was 13.76 ± 2.52. *Hprt1* mRNA expression was most stable across the passages, compared to *Gapdh* and *b-Act*. RefFinder software ranked the prospective housekeeping genes in order of the most to the least stable *as Hprt1, Gapdh and b-actin*.

The relative mRNA levels of these genes in harvested cochlea and vestibule membranous tissues are illustrated (Fig 9A). The targeted genes normalized mean Ct values when compared between cochlear and vestibular harvested tissues, demonstrated that m*yosin VIIa* expression levels in cochlear and vestibular tissues were almost identical (p = 0.947), whereas *prestin* expression was greater in cochlear tissue with a statistical significance at p = 0.029. Gene *Sox2* (p = 0.036) was significantly highly expressed in vestibular tissue whereas genes *nestin* (p = 0.113), *cytokeratin 18* (p = 0.12) and *vimentin* (p = 0.059) were not significantly different between porcine cochlear and vestibular harvested tissues.

**Fig 9.**
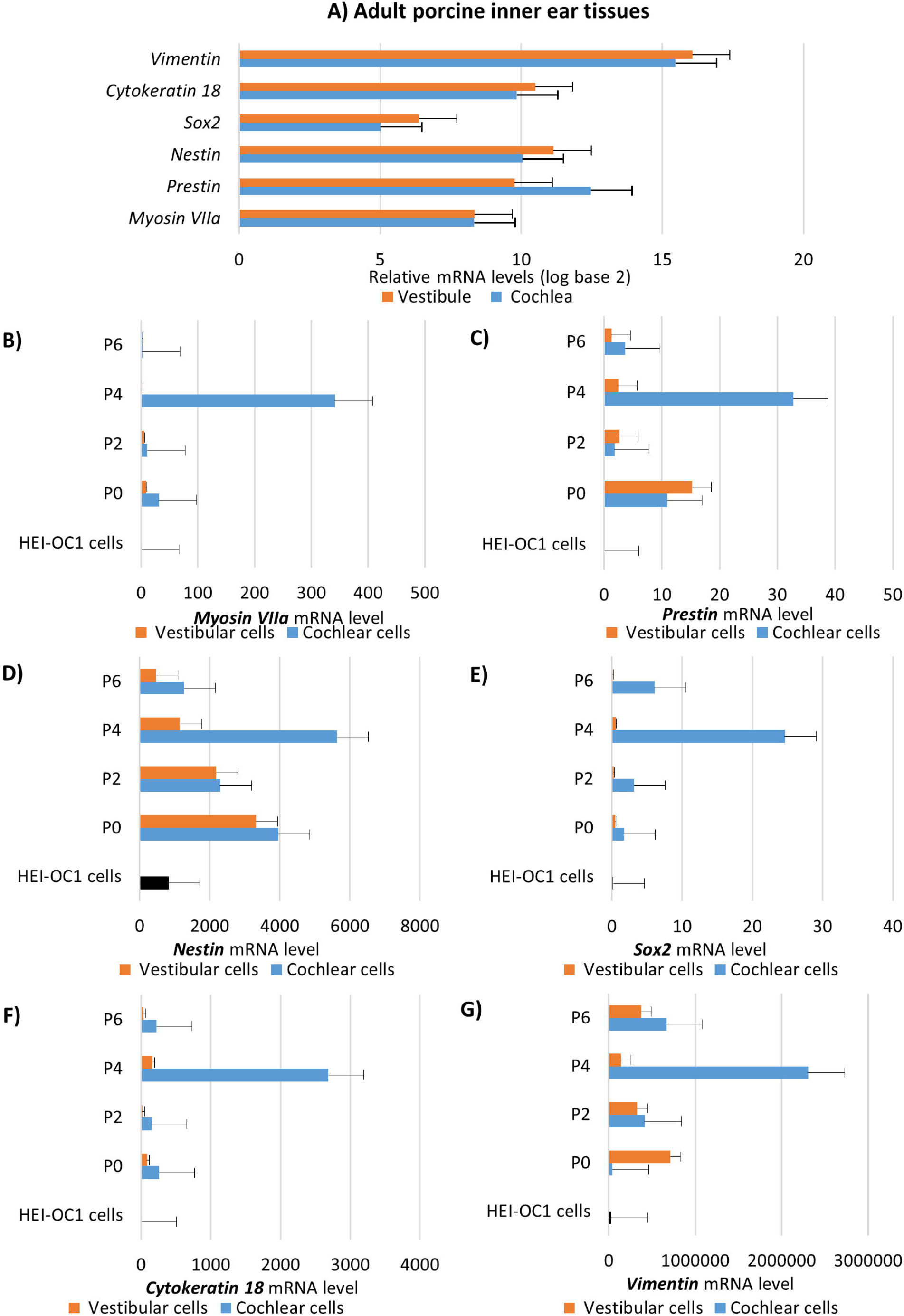
Relative gene expressions within cochlear (blue) and vestibular (orange) *in vitro* cell cultures. **A)** Relative mRNA levels of *myosin VIIa, prestin, nestin, Sox2, cytokeratin 18* and *vimentin* are presented for harvested adult porcine cochlea and vestibule in log base 2. **B)** to **G)** Relative mRNA levels of *myosin VIIa, prestin, nestin, Sox2, cytokeratin 18* and *vimentin* in *in vitro* cultures. Gene *hypoxanthine phosphoribosyltransferase 1* (*Hprt1*) was used as an endogeneous control. Positive control was HEI-OC1 cells (indicated by black). Number of replicates were three and the error bars represented the standard error.

The target genes’ normalized mean Ct values varied between porcine cochlear and vestibular cells. m*yosin VIIa* expression was significantly highly expressed at P4 (p = 0.000006) in cochlear compared with vestibular cells (Fig 9B). Similarly, *prestin* expression was significantly highly expressed at P4 (p = 0.00003) and at P6 (p = 0.024) in cochlear compared with vestibular cells (Fig 9C). Genes *Sox2* (Fig 9E) and *cytokeratin 18* (Fig 9F) were significantly highly expressed in cochlear cells at all tested passages: P0 (p = 0.017, p = 0.04), P2 (p = 0.001, p = 0.011), P4 (p = 0.00005, p = 0.00004) and P6 (p = 0.0001, p = 0.0002), respectively. *Nestin* at P2 (p = 0.015) and P4 (p = 0.01) (Fig 9D) and *vimentin* at P0 (p = 0.008) and P4 (p = 0.005) (Fig 9G) were also significantly highly expressed in cochlear compared with vestibular cells.

The normalized mean Ct values of all tested genes in porcine cochlear cells were significantly different (p < 0.05) across the passages 0 to 6 when compared with HEI-OC1 cells except for *nestin* expression at P6 (p = 0.26) and *vimentin* expression at P0 (p = 0.42) (Fig 9D and 9G, respectively). *Prestin* gene expression was undetectable in HEI-OC1 cells, thus statistical analysis was not performed.

The level of expression of of *cytokeratin 18*, and both HC markers *myosin VIIa* and *prestin* were significantly positively correlated [p = 0.005, Spearman correlation (τ) = 0.943] in porcine cochlear cultures, from harvested tissue to P6. Genes *myosin VIIa* and *prestin* expressions demonstrated a signifcant positive correlation in both porcine cochlear (p = 0.019, τ = 0.886) and vestibular (p = 0.005, τ = 0.943) cultures, from harvested tissue to P6. Porcine vestibular cultures also demonstrated a significant positive correlation between *nestin* and *prestin* mRNA levels from harvested tissue to P6 (p = 0.019, τ = 0.886).

## Discussion

We demonstrated that the pig’s labyrinth was anatomically similar to that of the human’s which is consistent with other studies [21, 22]. The maximal transverse and vertical axial diameters of the pig’s cochleas sampled are consistent with the findings reported in a large series of human cochleas [23]. Our use of an inserted cochlear implant electrode ascertained the length of the pig’s cochlea duct as 35mm in two specimens which is within the range of lengths of the human cochleas based on anatomical studies [24] and the mean lengths derived from CT data [25]; 28.0-40.1 mm and 34.62 mm, respectively. These findings indicate that the pig’s cochlear dimensions (length and cross sectional diameter) are similar to those of the human cochlea.

The development, function, and maintenance of inner ear sensory epithelia are heavily dependent upon the SCs [26], which are crucial in cochlea homeostasis. Hence, our aim was to develop an *in vitro* multi-cellular culture which mirrors the complexity of the human inner ear to facilitate the use of these cells in studies of hearing and balance disorders. Our findings indicate that primary cell cultures of adult porcine derived inner ear cochlear and vestibular cells are heterogeneous, highly adherent to plastic surfaces and demonstrate the presence of inner ear HC, SC and multipotent stem/progenitor cell characteristics based on SEM, fluorescence immunocytochemistry and RT-qPCR.

Sphere-forming cells were observed in both porcine cochlear and vestibular cell cultures and were similar to those observed in HEI-OC1 cultures. This suggests that these cultures contain cells with or that develop stem-like properties in the culture conditions we utilised. It remains unclear whether mammalian inner ear sphere-forming cells are endogenous stem cells or derived from SCs. In non-mammalian vertebrates such as birds, fish and amphibians, SCs are the most likely source of progenitor cells within the inner ear sensory epithelia [9]. These SCs can generate new HCs either via a regenerative response of dedifferentiation, proliferation and differentiation, or a direct phenotype conversion called trans-differentiation [9, 27]. It has been reported that vestibular sensory epithelia SCs display stem cell characteristics in mature guinea pigs and in adult humans *in vitro* [2, 3]. In mice, adult vestibular SCs replace damaged HCs to a certain extent [28], in contrast, cochlear SCs lose this capacity during the first few neonatal weeks as the inner ear completes its maturation [28-30]. Recent demonstrations of spontaneous HC regeneration in the immature neonatal mouse cochlea suggest that progenitor cells in that species are generated primarily by trans-differentiation of SCs and not by the proliferation of existing stem cells or dedifferentiation of SCs into stem cells [31, 32]. Although SCs with the capability for phenotypic conversion to HCs have been identified, such capability within the organ of Corti is only demonstrated in prenatal and neonatal rodent models where HCs and SCs are still developing [33-35]. Since the early postnatal cochlea of the mouse is immature, the relevance of these findings to regenerative therapies in adult humans is unclear [36].

In most previous studies, inner ear multipotent cells were induced to differentiate into cells expressing HC markers in part by adhesion to substrates, such as poly-D-lysine [37], poly-L-lysine [38, 28], fibronectin [39] and laminin [40]. Liu et al. [38] promoted differentiation of inner ear multipotent cells derived from postnatal day 0 mice cochlear sensory epithelia into functional HC-like cells containing characteristic stereocilia bundles through a two-step-induction method. Ding et al. [40] described similar techniques that induced the conversion of human embryonic stem cells into otic epithelial progenitors (OEPs) followed by the induced differentiation of OEPs into HC-like cells not only by substrate selection but also the deployment of conditioned media containing epidermal growth factor and all-trans retinoic acid. We did not use any growth factor media beyond DMEM plus 10% FBS and obtained HC-like cells from adult porcine inner ear cells which were similar on SEM to the HC-like cells obtained by Liu et al. [38] and Ding et al. [40]. Ding et al. [40] however demonstrated that HC-like cells that were induced on a substrate of mitotically inactivated chicken utricle stromal cells were functional and displayed more organized surface ciliary architecture than those that were induced on a poly-L-lysine substratum. They hypothesized that stromal substrate cells may release factors that are important for the development of functional HCs. Although, the HC-like cells that we generated displayed similar SEM characteristics, further work is required to determine their functional status. Importantly, we utilized the multiple cell types present in the inner ear as the starting point for our experiments thus we submit that any necessary factors that Ding et al. [40] hypothesized to be present in their utricle stomal cell substrate may be present in our *in vitro* cultures. Nonetheless these observations demonstrate a more direct method of generating HC-like cells from postmortem adult porcine labyrinth specimens than previously described techniques.

Immunofluorescence staining and RT-qPCR revealed that nestin was strongly expressed in both porcine cochlear and vestibular cultures across the passages 0 to 6 comparable to that seen in HEI-OC1 cell cultures. Nestin-positive cells have been found in tissue or organ-specific sites, where they serve as quiescent cells capable of proliferation, differentiation, and migration after their reactivation [41]. Nestin is associated with pluripotency in embryonic and induced pluripotent stem cells, as well as multipotency in spiral ganglion cells of the mature mouse [42]. Several studies have shown that nestin-positive cells derived from inner ear cells or embryonic stem cells serve as progenitors for sensory HC-like cells [43-45]. Chow et al. [42] found nestin-expressing cells adjacent to the inner HC layer in postnatal and young adult mice. Nestin-positive cells in the mature rat cochlea have been identified as SCs situated laterally, adjacent to outer HCs in the cochlea apex [46]. Lou et al.’s [47] study on cultivated cochlear cells derived from adult and neonatal mice suggests that stem/progenitor cells maintained their stemness but eventually lose the potential to differentiate into other cell types with age. Therefore, we propose that the strong nestin-expressing cells in porcine inner ear and HEI-OC1 cell cultures are multi-or oligo-potent progenitors that generate HC and SC-like inner ear cells especially when grown on poly-L-lysine coated surfaces. However, more work is required to identify the location and specific cell type(s) which demonstrate stem/progenitor cell activity within the porcine inner ear.

Mammalian stereocilia contain myosin VIIa which maintains the structural integrity of the HC bundles [48, 49]. HC-like cells with disorganized stereocilia were identified by SEM in the passaged porcine *in vitro* cultures. We also undertook SEM on cultured HEI-OC1 cells which to our knowledge has not been previoulsy reported and found HC-like with disorganized sterocilia similar to those in the porcine cultures. Although, myosin VIIa positive cells were identified using ICC in both porcine cochlear and vestibular cells from passages 0 through 6, relative *myosin VIIa* mRNA expression was identified at lower levels in passaged cultures than expected based on the ICC and SEM findings. The high number of HC-like cells on SEM and ICC may be due to the poly-L-lysine coating on the slides and coverslips used for these experiments as this substrate has been previously demonstrated to induce the conversion of multipotent stem/progenitor cells into HC-like cells [38, 40]. Similarly as the cell culture flasks used for qPCR were not coated with poly-L-lysine the same degree of induction of HC-like cells will not have occurred. Alternatively the differences in myosin VIIa cellular protein distribution and mRNA concentration may be a reflection of differences in the degree to which the cultured HC-like cells have differentiated to resemble functional HCs.

The co-localization of myosin VIIa and cytokeratin 18 in porcine cochlear and vestibular cells is consistent with that expected in immature HCs as cytokeratin is abundant in HC progenitor cells after which the level decreases progressively as HCs mature until it is no longer present in mature HCs [50]. The significant positive correleation between c*ytokeratin 18* and *myosin VIIa* mRNA expressions within the porcine cochlear cultures further supports the presence of immature HC-like cells across the passages.

The outer HC marker prestin belongs to the mammalian SLC26 family [51]. Recently, Park et al. [52] studied the HEI-OC1 auditory cells as a model for investigating prestin function. They confirm that under permissive conditions (33^°^C, 10% CO_2_), a condition in which HEI-OC1 cells proliferate *in vitro*, prestin is expressed mostly in the cytoplasm; in contrast, under non-permissive conditions (39^°^C, 5% CO_2_), a condition in which HEI-OC1 cells differentiate *in vitro*, total prestin expression is increased and localized to the plasma membrane. Our cell culture conditions (37^°^C, 5% CO_2_) are close to non-permissive conditions and we observed strong prestin expression in the plasma membranes of both adult porcine inner ear cells and HEI-OC1 cells *in vitro*. Similar to Adler et al. [53], we identified prestin protein in vestibular cell cultures; although prestin is primarily designated as an outer HC motor protein of the mammalian cochlea. The relative *prestin* mRNA level decreased from harvested tissues through P6 cell cultures and was undetectable in HEI-OC1 cells; however, plasma membrane prestin protein localization persisted across the passages when grown on coated 8-well chamber slides. These findings provide further evidence that the HC-like cells observed on SEM analysis of the porcine cell cultures grown on coated slides or cover slips possess differentiated HC-like characteristics.

The adult porcine derived inner ear cell *in vitro* cultures displayed a higher level of vimentin mRNA than the level in the harvested tissues. This may be a reflection of a high level of proliferation in the cultured cells as *vimentin* is associated with mitosis and cell growth [54]. Vimentin is present in several inner ear cell types which means that it lacks specificity as a marker of a single cell type. The cytoplasm of mammalian inner ear SCs including Deiters and inner pillar cells are vimentin rich [55] and in addition contain cytokeratin. Spiral ligament fibrocytes amongst others are a rich source of fibroblasts. In addition, the culture media contained FBS supplement a promoter of fibrobastic proliferation. Therefore the combination of a number of *vimentin* expressing cell types all of which are rapidly dividing may account for the increased *vimentin* mRNA levels in the cultured cells compared to the harvested tissues.

To our knowledge, housekeeping genes for RT-qPCR studies on porcine derived inner ear cells have not been previously published. We identified *Hprt1* as a suitable housekeeping gene for investigating genes in porcine inner ear tissue by comparison with *Gapdh* and *b-Act* which is consistent with Nygard et al.’s findings in other porcine tissues [56].

## Conclusion

Taken together, the similarity of the pig’s inner ear anatomy and cellular composition to that of humans suggest that the domestic pig can be considered as an animal model for the study of human inner ear disorders. Our findings suggest that adult porcine cochlea and vestibule tissue have the capacity to form new HC populations. Furthermore, we found evidence for multipotent stem/progenitor cells in adult derived inner ear *in vitro* cultures though additional work is required to identify the cell type(s) and the location of these cells within the porcine inner ear.

## Acknowledgements

The authors thank Dr. F. Kalinec, Laboratory Auditory Cell Biology, Dept. of Head & Neck Surgery, David Geffen Scholl of Medicine at UCLA, Los Angeles, CA, USA for gifting HEI-OC1 cells. Our sincere thanks to Mr. Derrick Horne, Technical Specialist, UBC Bioimaging Facility, Vancouver Campus for his technical assistance to carry out the scanning electron microscopy on passaged in inner ear cell *in vitro* cultures.

## Author contributions

D.A.N. proposed the study and obtained funding. D.A.N., A.S., T.A.C. and P.W. designed the experiments and interpreted the results. P.W. and A.S. drafted the paper. D.A.N. and E.H. revised the paper. A.S., T.A.C., P.W., B.Z., E.H., G.H. and J.K. developed the cell cultures.

P.W. performed the cellular characterization studies. P.W. analysed the data. A.S. and T.A.C. performed the anatomical studies. All authors reviewed the final manuscript.

