## Supplementary material for "Adult porcine (*Sus scrofa*) derived inner ear cells possessing multipotent stem/progenitor cell characteristics in *in vitro* cultures": S1 Table

| **Genes**  ***(Sus scrofa)*** | **Primers** | **Sequences 5’ - 3’** |
| --- | --- | --- |
| *Myosin VIIa* | Forward | CCAAGGACATCCTGACCACT |
|  | Reverse | CCACCTTCCCACTGTTCACT |
| *Prestin* | Forward | TCCTTGTCTCGAAGCCTTGT |
|  | Reverse | ACATCCCTTTCAGGTTGACG |
| *Nestin* | Forward | CTTCCAAGACCCTGAAGAGA |
|  | Reverse | TTCCAACACGGGCTCTAT |
| *Sox2* | Forward | CATGCTGATCATGTCCCGTAGGT |
|  | Reverse | GCCCTGCAGTACAACTCCATG |
| *Cytokeratin 18* | Forward | ACCTCAGGACCTCAGCAAGA |
|  | Reverse | CTCATGGAGTCCAGGTCGAT |
| *Vimentin* | Forward | CATCAACACCGAGTTCAAGA |
|  | Reverse | GCACCTTGTCGATGTAGTT |
| *Gapdh* | Forward | GTCGGTTGTGGATCTGACCT |
|  | Reverse | AGCTTGACGAAGTGGTCGTT |
| *b-Act* | Forward | CACGCCATCCTGCGTCTGGA |
|  | Reverse | AGCACCGTGTTGGCGTAGAG |
| *Hprt1* | Forward | GGACTTGAATCATGTTTGTG |
|  | Reverse | CAGATGTTTCCAAACTCAAC |

**S1 Table. Primer sequences used for adult porcine derived inner ear samples.** Forward and reverse primers are shown for eight genes expressed in inner ear hair cells and supporting cells of the cochlea and vestibular cells. *Gapdh-Glyceraldehyde-3-phosphate dehydrogenase; b-Act- beta-actin; Hprt1- hypoxanthine phosphoribosyltransferase 1*

| **Genes**  **(*Mus musculus*)** | **Primers** | **Sequences 5’ - 3’** |
| --- | --- | --- |
| *Myosin 7a* | Forward | CAACATGAAACGCAACAACC |
|  | Reverse | CCAAAGCGGCTAGAGTTGTC |
| *Prestin* | Forward | ACAGTGTGGATGTCGTTGGA |
|  | Reverse | CCATGCTTATTTGCCAAGGT |
| *Nestin* | Forward | CCAGAGCTGGACTGGAACTC |
|  | Reverse | ACCTGCCTCTTTTGGTTCCT |
| *Sox2* | Forward | AAGGGTTCTTGCTGGGTTTT |
|  | Reverse | AGACCACGAAAACGGTCTTG |
| *Cytokeratin 18* | Forward | AGACTTGGTGGTGACAACTGTGG |
|  | Reverse | ATCGAGGCACTCAAGGAAGA |
| *Vimentin* | Forward | CGCAGCCTCTATTCCTCATC |
|  | Reverse | GTAGTTGGCAAAGCGGTCAT |
| *Hprt1* | Forward | GCCCCAAAATGGTTAAGGTT |
|  | Reverse | TTGCGCTCATCTTAGGCTTT |

**Supplementary table 2: Primer sequences used to amplify the target genes in HEI-OC1 cells.** *Sox2- SRY (sex determining region Y-box2); Hprt1- hypoxanthine guanine phosphoribosyl transferase 1.*
